# The DEAD-box helicase RCF1 plays roles in miRNA biogenesis and RNA splicing in *Arabidopsis*

**DOI:** 10.1101/2023.04.03.535409

**Authors:** Chi Xu, Zhanhui Zhang, Juan He, Yongsheng Bai, Lin Liu, Jihua Tang, Guiliang Tang, Xuemei Chen, Beixin Mo

## Abstract

RCF1 is a highly conserved DEAD-box helicase found in yeast, plants and mammals. Studies about the functions of RCF1 in plants are limited. Here we uncovered the functions of RCF1 in *Arabidopsis thaliana* as a player in pri-miRNA processing and splicing, as well as in pre-mRNA splicing. A mutant with miRNA biogenesis defects was isolated and the defect was traced to a recessive point mutation in *RCF1* (*rcf1-4*). We show that RCF1 promotes D-body formation and facilitates the interaction between pri-miRNAs and HYL1. Finally, we show that intron-containing pri-miRNAs and pre-mRNAs exhibit a global splicing defect in *rcf1-4*. Together, this work uncovers roles for RCF1 in miRNA biogenesis and RNA splicing in *Arabidopsis*.

**One-sentence summary:** RCF1 promotes not only the processing of pri-miRNAs, but also the splicing of intron-containing pri-miRNAs, therefore promotes miRNA biogenesis.

## Introduction

MicroRNAs (miRNAs), which are composed of 20∼24 nucleotides, negatively regulate gene expression by targeting complemental sequences (Rogers and Chen, 2013; Song et al., 2019; Li and Yu, 2021; Zhang et al., 2022). Similar as pre-mRNAs, MIRNA (MIR) transcripts also undergo 5’ cap addition, 3’ polyadenylation, and splicing (for intron-containing transcripts). (Rogers and Chen, 2013; Hunt, 2014; Ramanathan et al., 2016; Deng and Cao, 2017). The pri-miRNAs with typical stem-loop structures are processed to produce pre-miRNAs, which are further processed to miRNA/miRNA* duplexes by Dicer-like 1(DCL1), a RNAse III endoribonuclease, (Kurihara and Watanabe, 2004). HYPONASTIC LEAVES1 (HYL1) and SERRATE (SE) are the two cofactors of DCL1 in D-bodies by increasing the the accuracy and efficiency of miRNA processing (Kurihara et al., 2006; Lobbes et al., 2006; Dong et al., 2008). D-bodies are nuclear foci containing the microprocessor components and are thought to be sites of pri-miRNA processing (Fang and Spector, 2007; Zhu et al., 2013; Xie et al., 2021), but processing also takes place elsewhere, such as on chromatin (Stepien et al., 2022). After cleavage, the miRNA/miRNA* duplexes undergo HUA ENHANCER 1 (HEN1)-mediated methylation, which maintains their stability (Yu et al., 2005). Then, miRNAs are loaded into ARGONAUTE1 (AGO1) protein and form miRNA-induced silencing complexes (miRISCs) (Baumberger and Baulcombe, 2005; Qi et al., 2005; Bologna et al., 2018). miRISCs repress their target mRNAs mainly by promoting mRNA degradation or/and translational repression. In recent years, dozens of regulatory factors have been identified to affect pri-miRNAs’ structure, splicing, stability, and loading to the microprocessor, thus impacting miRNA biogenesis(Li and Yu, 2021; Zhang et al., 2022).. For example, Chromatin remodelling factors2 (CHR2) can remodel the structure of pri-miRNAs (Wang et al., 2018); the stability of pri-miRNAs is regulated by Protein Pleiotropic Regulatory Locus 1(PRL1, a WD-40 protein),(Zhang et al., 2014), MOS4-associated complex 3 (MAC3, a U-box type E3 ubiquitin ligase) (Li et al., 2018), MOS4-associated complex 5 (MAC5A)(Li et al., 2020) and AAR2 (Fan et al., 2022); Glycine-rich RNA-binding protein 7 (GRP7) (Koster et al., 2014), STABILIZED1 (STA1) (Ben et al., 2013), Increased Level of Polyploidy1-1D (ILP1)/NTC-related protein 1(NTR1) (Wang et al., 2019), and Serrate-Associated Protein 1 (SEAP1) (Li et al., 2021) are important for pri-miRNA splicing; the loading of pri-miRNAs to the microprocessor is regulated by Modifier of snc1;2 (MOS2) (Wu et al., 2013), Debranching RNA Lariats 1 (DBR1) (Li et al., 2016), and Short Valve 1 (STV1) (Li et al., 2017). Apart from the core components of the microprocessor, many other proteins participate in miRNA biogenesis by interacting with the core factors, such as MOS4-associated Complex 7 (MAC7) (Jia et al., 2017), AAR2 (a homolog of a U5 snRNP assembly factor) (Fan et al., 2022), CBF gene expression 3 (RCF3) (Karlsson et al., 2015), CHR2 (Wang et al., 2018), MOS4-associated Complex 5 (MAC5) (Li et al., 2020), TOUGH (TGH) (Ren et al., 2012), Cell Division Cycle 5(CDC5) (Zhang et al., 2013), Receptor of Activated C Kinase 1 (RACK1) (Speth et al., 2013), STV1(Li et al., 2017), and SEAP1 (Li et al., 2021).

The DEAD-box RNA helicase family exists in all eukaryotes and most prokaryotes, and is the largest family of RNA helicases (Nidumukkala et al., 2019; Wang et al., 2020). In plants, studies revealed DEAD-box RNA helicases to be involved in various processes in gene expression, including chromatin remodeling, transcription, ribosome biogenesis, pre-mRNA splicing, RNA degradation, translation, and nucleocytoplasmic transport (Ehrnsberger et al., 2019; Kesarwani et al., 2019; Capel et al., 2020; Lu et al., 2020; Takagi et al., 2020; He et al., 2021; Tyagi et al., 2021). The *Arabidopsis* genome has 58 genes encoding DEAD-box RNA helicases, named as AtRH1-58 (Mingam et al., 2004), of which, AtRH6/8/12 have been found to affect microRNA (miRNA) processing through interacting with SE (Li et al., 2021), while AtRH27 was found to associate with pri-miRNAs and interact with HYL1 and SE (Hou et al., 2021). In this work, a mutation in *Arabidopsis RCF1* (encoding a DEAD-box helicase) was isolated using a genetic screen for genes involved in miRNA biogenesis. We found that RCF1 acts in miRNA biogenesis by enhancing D-body formation, promoting the interaction between HYL1 and pri-miRNAs and facilitating the splicing of intron-containing pri-miRNAs. These findings provide insights into the post-transcriptional regulation of miRNA biogenesis.

## Results

### A miRNA biogenesis defective mutant was isolated by EMS mutagenesis

The *amiR-SUL* transgenic line is a visual reporter of miRNA activity and a good genetic material for the study of miRNA biogenesis (de Felippes et al., 2011). In the *amiR-SUL* line, using an artificial miRNA (amiR-SUL) to target *SUL* (also known as *CHLORINA42*), driven by a phloem specific promoter (*SUC2*), leads to vein-centered leaf albinoing (de Felippes et al., 2011). From a mutant population of amiR-SUL induced by ethyl methanesulfonate (EMS), we have reported a series of miRNA pathway genes, including *PP4R3A* (Wang et al., 2019), *CPR1* and *ABA1* (Cai et al., 2018), and *RBV* (Liang et al., 2022). Here, we obtained a suppressor mutant *sar2450 amiR-SUL* with reduced leaf albinoing (Figure 1A and B; Supplemental Figure S1A). The reduced leaf albinoing phenotype suggested that *sar2450 amiR-SUL* has a higher SUL protein level. Western blotting confirmed increased SUL protein accumulation in the *sar2450 amiR-SUL* (Supplemental Figure S1 B). Besides reduced leaf albinoing, *sar2450 amiR-SUL* also displayed pleiotropic developmental phenotypes including later flowering, unusual inflorescences, increased petal number, shorter silique length (Figure 1 A to I), some of which are reminiscent of mutants defective in miRNA biogenesis.

**Figure 1.**
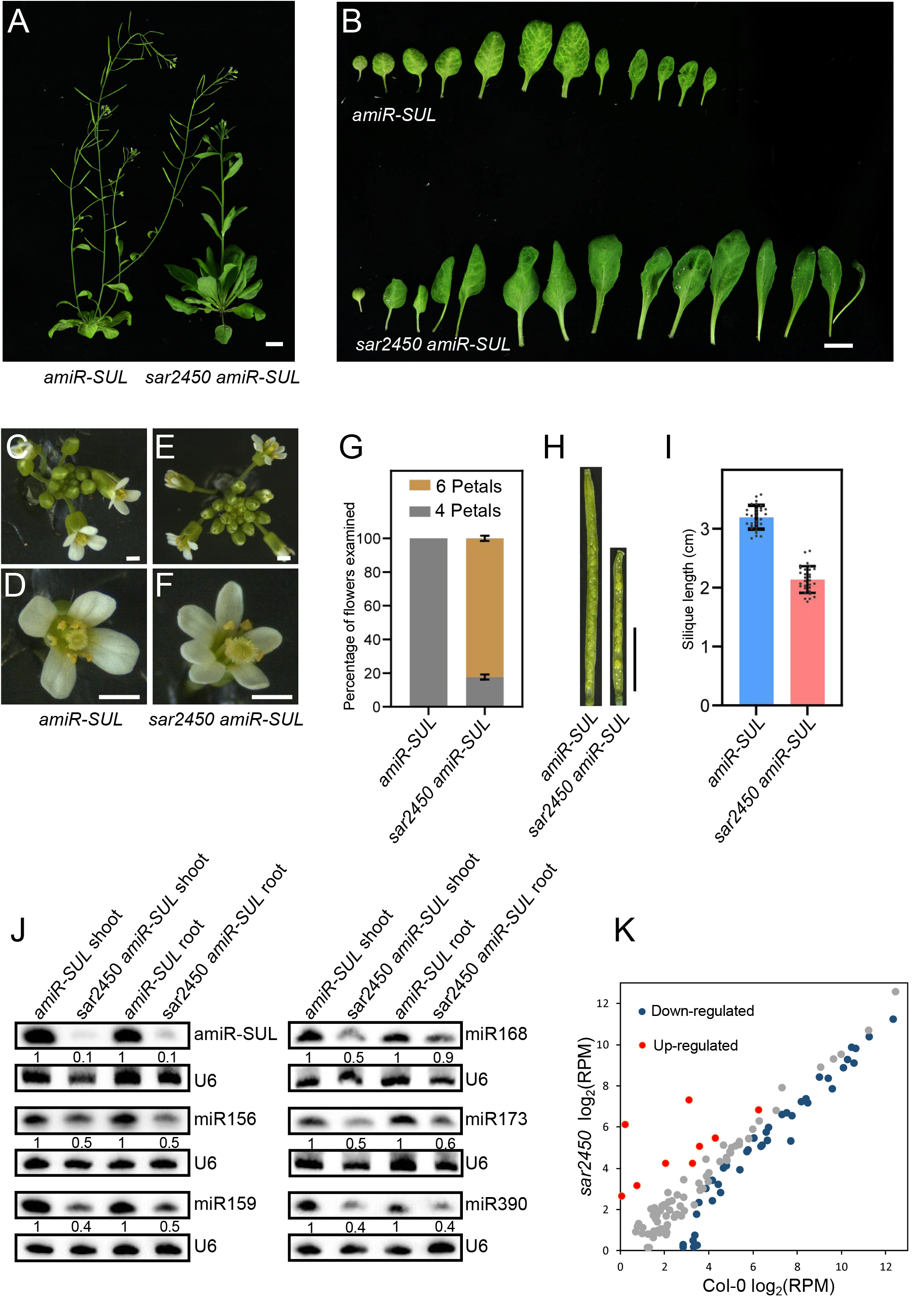
Isolation of a miRNA biogenesis suppressor *sar2450 amiR-SUL* from an *amiR-SUL* line with pleiotropic developmental defects and a global reduction in miRNA accumulation. (A) Morphological phenotypes of 45-day-old *amiR-SUL* and *sar2450 amiR-SUL* plants. Bar = 1 cm. (B) Images of rosette leaves from 45-day-old *amiR-SUL* and *sar2450 amiR-SUL* plants. Bar = 1 cm. (C-G) Images of the aberrant flowers and inflorescences in *sar2450 amiR-SUL*. Bar = 1 cm. (H) Images of the mature siliques showing the reduced silique length in *sar2450 amiR-SUL*. (I) Measurements of silique length indicated that *sar2450 amiR-SUL* had significantly shorter siliques. Error bars indicate SD (standard deviation), (n = 10 siliques). (J) Northern blotting analysis showed reduced levels of artificial amiR-SUL and endogenous miRNAs in *sar2450 amiR-SUL.* U6 RNA served as a loading control. The numbers below the gel images represent the relative amount. (K) A scatter plot showing the accumulation of miRNAs in *amiR-SUL* and *sar2450 amiR-SUL*. The miRNA abundance were normalized by calculated reads per million (RPM). The red dots indicate up-regulated miRNAs in *sar2450 amiR-SUL*, and the blue dots indicate down-regulated miRNAs in *sar2450 amiR-SUL*.(*P < 0.05).

We analyzed miRNA levels in the shoots and roots of *amiR-SUL* and *sar2450 amiR-SUL* by northern blotting. Results showed that *sar2450 amiR-SUL* plants exhibited significantly reduced levels of amiR-SUL as well as all examined endogenous miRNAs, such as miR156, miR159, miR168, miR173 and miR390 (Figure 1 J). Correspondingly, target genes of these miRNAs displayed up-regulated expression in *sar2450 amiR-SUL* mutants (Supplemental Figure S1C). These results indicated a general defect in miRNA accumulation in the *sar2450 amiR-SUL*.

We obtained the *sar2450* mutant without the *amiR-SUL* transgene by crossing *sar2450 amiR-SUL* with Col-0. The mutant *sar2450* showed a similar phenotype to that of *sar2450 amiR-SUL* (Supplemental Figure S1 A). Western blotting revealed that SUL protein levels did not differ significantly in *sar2450* compared to Col-0 (Supplemental Figure S1B). We performed sRNA-seq in Col-0 and *sar2450* seedings for a genome-wide analysis of miRNA accumulation. Each genotype had three biological replicates, which were highly correlated by PCA (Principal Component Analysis) analysis (Supplemental Figure S2 A). Accumulation of miRNAs showed a global reduction in *sar2450* (Supplemental Figure S2 B). 35 miRNAs were significantly down-regulated and 8 miRNAs were significantly up-regulated in the mutant (Figure 1K; Supplemental Dataset S1). These downregulated miRNAs did not differ significantly in the orientation (base-to-loop or loop-to-base) of pri-miRNA processing, or the length of pri-miRNAs compared to the upregulated or unaffected miRNAs (Supplemental Figure S3 A, B). Since the production of ta-siRNAs is triggered by miRNAs, we analyzed the reads of ta-siRNAs generated from *TAS2* and *TAS3A* transcripts. The accumulation of these ta-siRNAs was significantly decreased in *sar2450* (Supplemental Figure S4 A and B), indicating that ta-siRNA biogenesis was impaired indirectly in *sar2450*.

### The phenotype of *sar2450 amiR-SUL* was caused by a mutation in *RCF1*

In order to pinpoint the *sar2450* mutation, we back-crossed *sar2450 amiR-SUL* with the parental line *amiR-SUL*. All the BC_1_F_1_ individuals showed the *amiR-SUL* phenotype, indicating that *sar2450* was a recessive mutation. In a total of 1066 BC_1_F_2_ (BC_1_F_1_ inbred progenies) plants, 256 (24%) exhibited *sar2450 amiR-SUL* phenotypes, suggesting that the *sar2450 amiR-SUL* phenotype was caused by a single, nuclear mutation. The pooled DNA from the BC_1_F_2_ plants exhibiting the *sar2450 amiR-SUL* phenotype was used for re-sequencing. A G1732-to-A mutation in *AT1G20920* was identified in *sar2450* that converted the corresponding amino acid from glycine to glutamic acid (Figure 2A). *AT1G20920* encodes a DEAD-box RNA helicase known as RCF1, which contains an RD/RS domain, a DPLP motif, and 9 helicase motifs (motif Q, I, Ia, Ib, II, III, IV, V, and VI) (Figure 2A). We named this mutant as *rcf1-4*, because three mutants (*rcf1-1*, *rcf1-2* and *rcf1-3*) had previously been reported; *rcf1-2* and *rcf1-3* are T-DNA insertion mutants with the inserted DNA being located in the 3’ UTR of *RCF1* (Guan et al., 2013). *rcf1-1* is a point mutation, which results in an amino acid change from glycine to tyrosine between domains VI and V (Figure 2 A) (Guan et al., 2013). *rcf1-1* in the L*er* background and *rcf1-4* in the *Col-0* background had similar phenotypes, such as abnormal inflorescences and shorter silique length (Supplemental Figure S5 A to C). Northern blotting analysis showed that the accumulation of miRNAs in *rcf1-1* was also reduced as in *rcf1-4* (Supplemental Figure S5 D). These results further confirmed that *RCF1* was required for miRNA biogenesis. To determine whether the *rcf1-4* mutation contributed to the mutant phenotypes of *rcf1-4 amiR-SUL*, a construct carrying the coding region of *RCF1* fused with *GFP* driven by an *RCF1* promoter was generated and introduced into *rcf1-4 amiR-SUL* (Figure 2 B). The *pRCF1:RCF1-GFP* transgene fully rescued the phenotypes of *rcf1-4 amiR-SUL*, such as leaf color, SUL protein abundance, and miRNA levels (Figure 2B to D). Similarly, *pRCF1:RCF1-GFP1* also complemented the *rcf1-4* morphological phenotypes in the Col-0 background (Supplemental Figure S6 A to D). Northern blotting analysis revealed that the decreased accumulation of two ta-siRNAs, siR255 and 5’D8, as well as their triggers (miR173 and miR390) in *rcf1-4* was restored by *pRCF1:RCF1-GFP1* (Supplemental Figure S7 A and B). Moreover, siR255 and 5’D8 were also decreased in the *rcf1-1* mutant (Supplemental Figure S7 C and D), further confirming that RCF1 is required for ta-siRNA and miRNA biogenesis.

**Figure 2.**
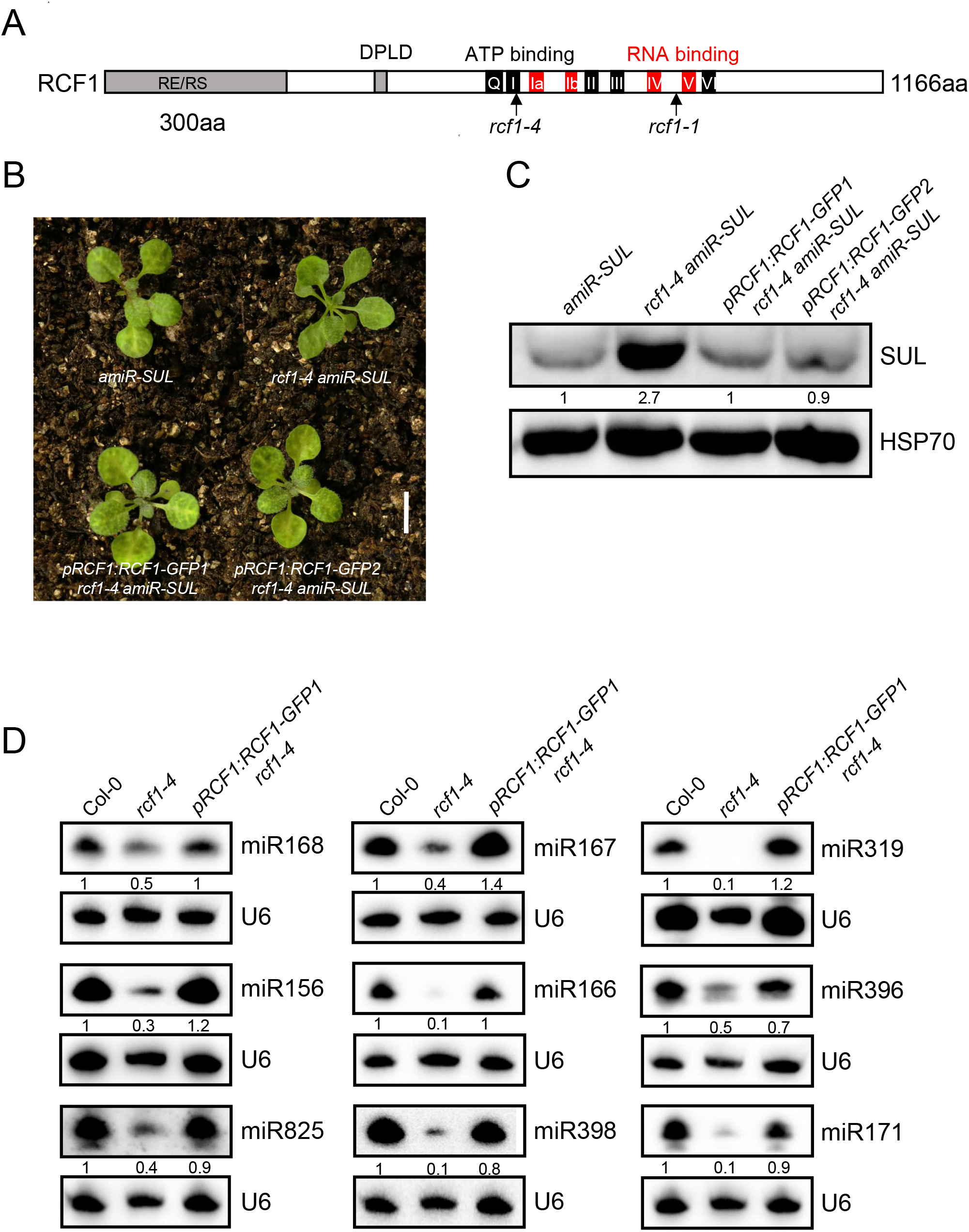
A mutation in *RCF1* accounts for the phenotypes of *sar2450*. (A) Diagram of RCF1 protein showing its domain structures. The boxes in the diagram represent the RD/RS domain, the DPLP motif, and helicase motifs Q, I, Ia, Ib, II, III, IV, V, and VI, respectively. The regions represented by the black rectangles contribute to ATP binding, and the regions represented by the red rectangles contribute to RNA binding. aa, amino acids. (B) Images of 14-day-old seedlings of the genotypes as indicated below. (C) Western blotting analysis of the protein accumulation of SUL in the genotypes as indicated. HSP70 was used as a loading control. (D) Northern blotting analysis of endogenous miRNAs in Col-0, *rcf1-4* and *pRCF1:RCF1-GFP1 rcf1-4*. U6 RNA served as a loading control. The numbers below the gel images represent the relative amount.

### RCF1 does not affect *MIR* transcription

*MIR* genes are first transcribed into pri-miRNAs, which undergo multiple processing steps to produce mature miRNAs. To determine the role of RCF1 in *MIR* transcription, we performed RT-qPCR analysis for pri-miRNAs in *rcf1-4* and wild type Arabidopsis (Figure 3 A). We found that all examined pri-miRNAs showed higher levels of accumulation in *rcf1-4*. To determine the effects of *RCF1* on *MIR* promoter activities, we crossed a *pMIR167a*:*GUS* line (Liang et al., 2022) with *rcf1-4*. GUS staining indicated that *MIR* promoter activity was unaffected in *pMIR167a:GUS rcf1-4* plants compared to *pMIR167a*:*GUS* plants (Figure 3 B). RT-qPCR analysis showed similar levels of *GUS* transcripts in these two genotypes, consistent with the GUS staining results (Figure 3 C). These results suggested that *RCF1* does not affect *MIR* gene transcription. The over accumulation of pri-miRNAs could be a result of compromised pri-miRNA processing, as observed in *dcl1*, *hyl1*, and se mutants (Kurihara and Watanabe, 2004; Kurihara et al., 2006; Lobbes et al., 2006).

**Figure 3.**
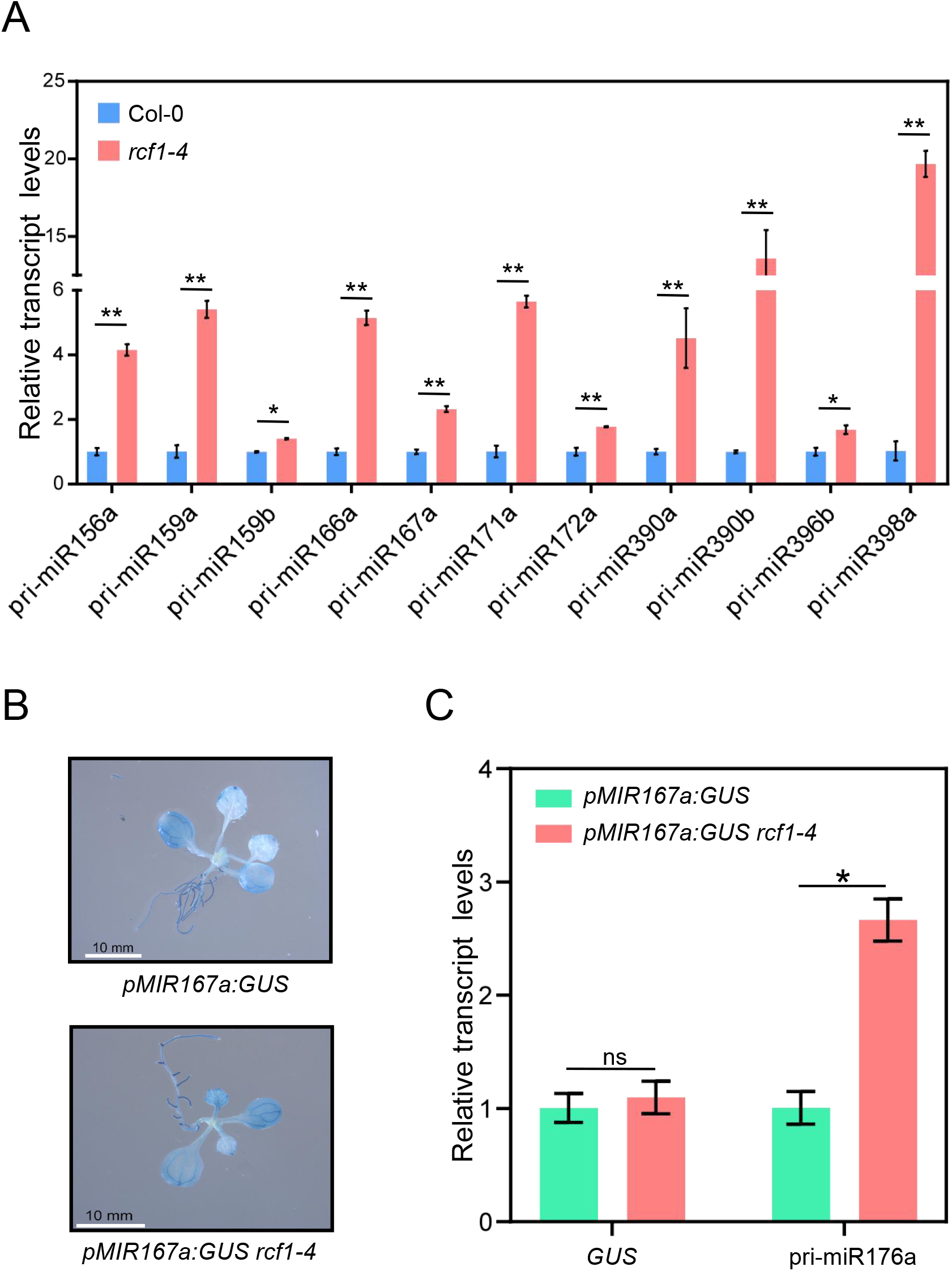
RCF1 promotes pri-miRNA accumulation without affecting *MIR* promoter activities. (A) RT-qPCR analysis of the accumulation of pri-miRNAs in Col-0 and *rcf1-4*. *UBQ5* was used as the internal control. (Error bars represent SD(standard deviation) of three replicates, *P < 0.05). (B) Representative images of GUS staining of *pMIR167a:GUS* and *pMIR167a:GUS rcf1-4* transgenic plants. Bars = 1 mm. Levels of the *GUS* transcript and pri-miR167a in *pMIR167a::GUS* and *pMIR167a:GUS rcf1-4* plants as determined by RT-qPCR. (Error bars represent SD (standard deviation) of three replicates, *P < 0.05).

### RCF1 promotes D-body formation

D-body may be a site of pri-miRNA processing (Fang and Spector, 2007; Zhang et al., 2020a; Xie et al., 2021). To determine whether RCF1 affects D-body formation, we crossed the D-body marker line *pHYL1:HYL1-YFP* with *rcf1-4*. The distribution of HYL1-YFP in *pHYL1:HYL1-YFP rcf1-4* and *pHYL1:HYL1-YFP* root meristem cells were observed by confocal microscopy (Figure 4 A). The number of HYL1-YFP-labeled D-bodies was significantly reduced in *pHYL1:HYL1-YFP rcf1-4* compared to that in *pHYL1:HYL1-YFP* (Figure 4 B), indicating that RCF1 may affect the processing of pri-miRNAs by promoting D-body formation or localization of HYL1 to D-bodies. RCF1-GFP is localized in the nucleus (Guan et al., 2013), which was similar to that of SE-mRuby3 (Liang et al., 2022). However, we did not detect any interactions between RCF1 and SE through yeast two-hybrid or Co-IP analysis. The levels of DCL1, HYL1 and SE proteins were higher in the *rcf1-4* mutant than wild type (Figure 4 C), indicating that the defect in miRNA biogenesis in *rcf1-4* was not due to reduced accumulation of microprocessor core proteins. Considering that HYL1 is a nucleo-cytoplasmic shuttling protein, reduced HYL1 protein distribution in the nucleus may also cause defects in miRNA biogenesis. We extracted nuclei from *rcf1-4* and Col-0, and western blotting revealed that the protein levels of HYL1, as well as DCL1 and SE, were significantly increased in the nucleus in *rcf1-4* (Figure 4 D). Taken together, these results showed that the defects in miRNA biogenesis or D-body formation in *rcf1-4* were not due to reduced levels of pri-miRNAs or microprocessor core proteins and suggested that *RCF1* might enhance miRNA biogenesis through its role in D-body formation.

**Figure 4.**
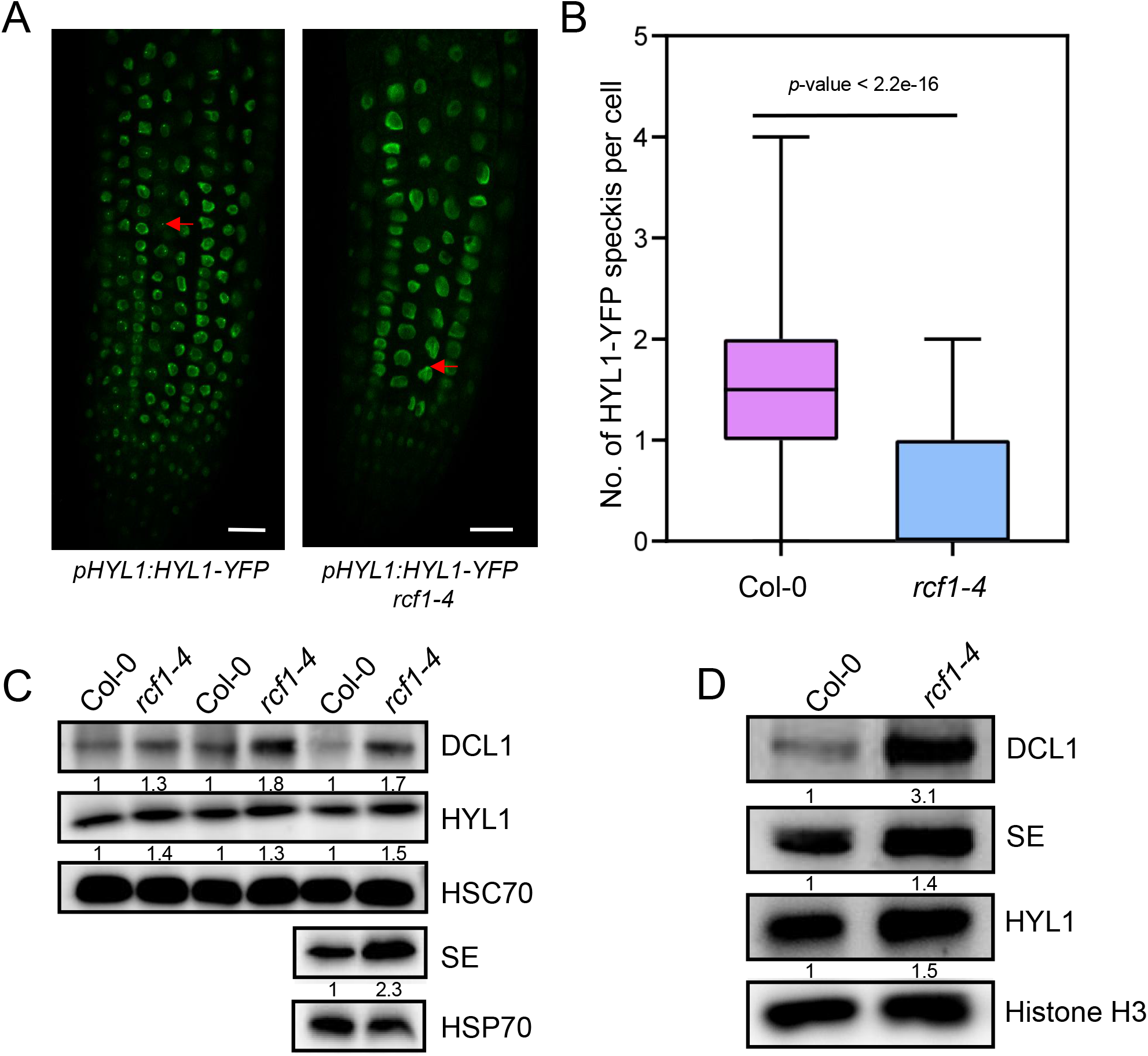
RCF1 facilitates the proper localization of HYL1 in D-bodies. (A) Representative images of HYL1-YFP signals in root cells of 2-week-old seedlings of *pHYL1:HYL1-YFP* and *pHYL1:HYL1-YFP rcf1-4*. Arrows pointed out D-bodies. Bars = 5 μm. (B) Number of D-bodies per cell in Col-0 and *rcf1-4*. 200 cells were detected for each genotype. (C) Western blotting analysis of the protein levels of microprocessor components in Col-0 and *rcf1-4* seedlings. HSP70 served as a loading control. (D) Western blotting analysis of microprocessor components in the nuclear fraction in Col-0 and *rcf1-4* seedlings, Histone H3 served as a loading control.

### RCF1 is required for the binding of pri-miRNAs by HYL1

HYL1 is known to bind pri-miRNAs and assist DCL1 in pri-miRNA processing (Hiraguri et al., 2005; Kurihara et al., 2006; Curtin et al., 2008; Dong et al., 2008; Wu et al., 2013). We examined the interaction between HYL1 and pri-miRNAs by RNA immunoprecipitation (RIP) using HYL1 antibodies, using the amounts of HYL1 protein by IP as the loading control (Figure 5 A). Following HYL1 IP, pri-miRNAs associated with HYL1 were detected by RT-qPCR, and results showed that all examined pri-miRNAs (pri-miR156a, pri-miR159a, pri-miR167a, pri-miR172a, pri-miR396a) were significantly reduced in *rcf1-4* (Figure 5 B to F), indicating that RCF1 promotes the binding of pri-miRNAs by HYL1.

**Figure 5.**
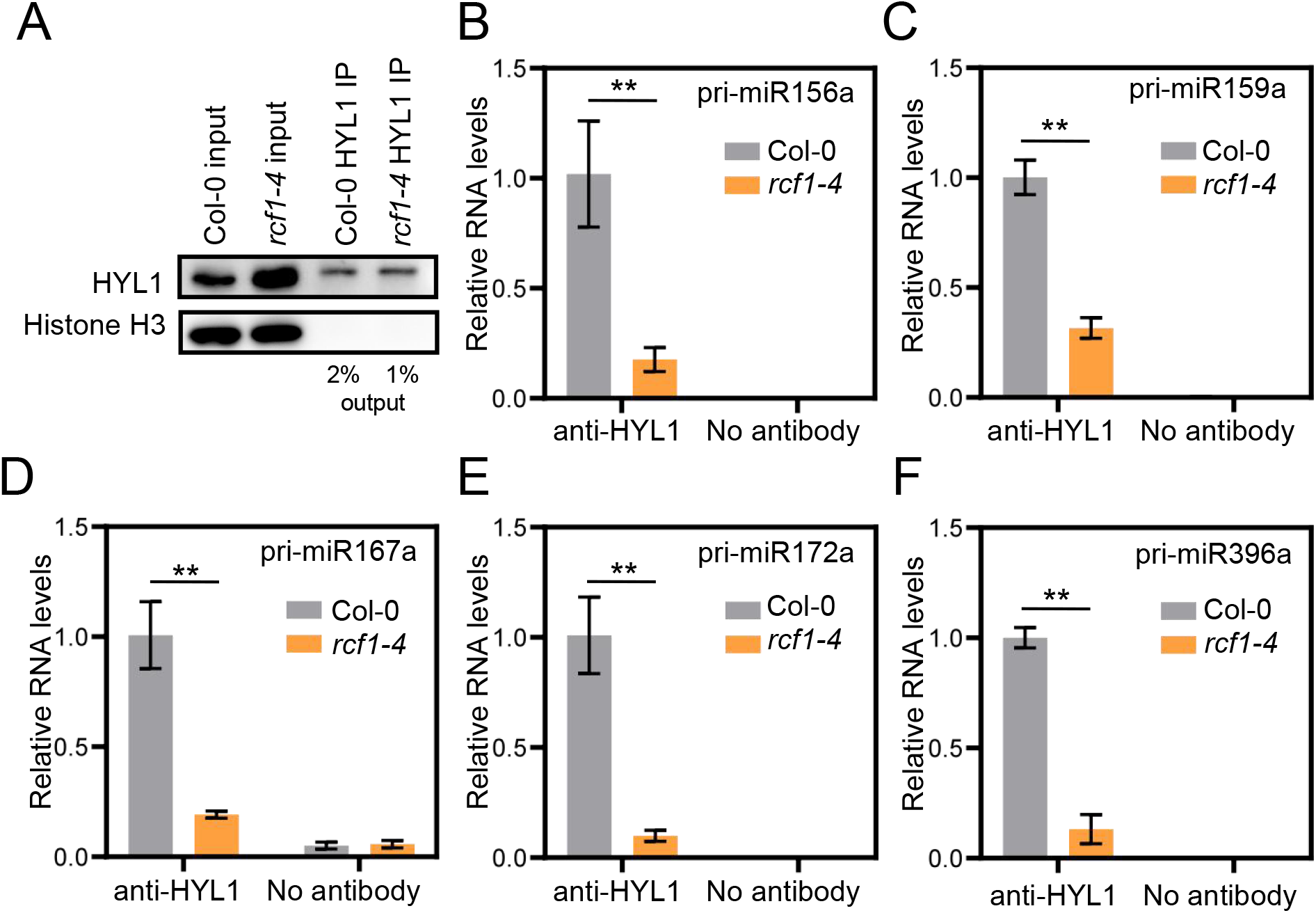
RCF1 facilitates the binding of pri-miRNAs by HYL1. (A) Western blotting analysis of HYL1 and histone H3 after HYL1 IP in Col-0 and *rcf1-4* seedlings. For the input samples, Histone H3 was used as a loading control. For the IP samples, Western blotting analysis to quantify HYL1 protein levels. (B-F) HYL1-bound pri-miRNAs detected by RT-qPCR. RNAs from the same amount of IP-ed HYL1 were used for RT-qPCR. (Error bars represent SD (standard deviation) of three replicates, **P < 0.01).

### RCF1 promotes pri-miRNA splicing

A large number of *MIRs* in Arabidopsis contain introns or are located in introns of host genes (intronic miRNAs) (Figure 6 A) (Yang et al., 2012; Stepien et al., 2017). For the pri-miRNAs containing introns or located in introns, intron splicing is a non-negligible factor for their subsequent processing (Bielewicz et al., 2013; Jia and Rock, 2013; Schwab et al., 2013; Knop et al., 2017; Stepien et al., 2017). On the one hand, the splicing of introns directly affects the processing efficiency of pri-miRNAs, such as pri-miR161, pri-miR163 and pri-miR172b. On the other hand, for pri-miRNAs located within introns of coding genes or non-coding genes, the introns need to be spliced from the genes before they can be efficiently processed, such as *MIR402*. Previous studies have shown that some pre-mRNA splicing factors promote miRNA biogenesis, such as CBP20 and CBP80(Kim et al., 2008), STA1 (Ben et al., 2013), SMA1 (Li et al., 2018), MAC3A/MAC3B (Li et al., 2018) and PRL1/PRL2 (Zhang et al., 2014), SEAP1 (Li et al., 2021). The accumulation of most miRNAs was significantly reduced in *rcf1-4*, regardless of whether their precursors contained introns or not (Figure 6 A-B). However, those miRNAs whose precursors contained introns decreased more significantly than the miRNAs whose precursors did not contain introns (Figure 6 B). Interestingly, the accumulation of intronic miRNAs was unaffected in *rcf1-4* compared to wild type (Figure 6 B). We speculated that besides promoting D-body formation and pri-miRNA’s association with HYL1, RCF1 may also enhance the biogenesis of intron-containing miRNAs by enabling the splicing of their precursors. We further performed RT-PCR to examine the splicing of some intron-containing pri-miRNAs (pri-miR156a, pri-miR156c, pri-miR172a) with intron-flanking primers. All the tested pri-miRNAs displayed intron retention defects in *rcf1-4* (Figure 6 C), confirming that RCF1 affects pri-miRNA splicing.

**Figure 6.**
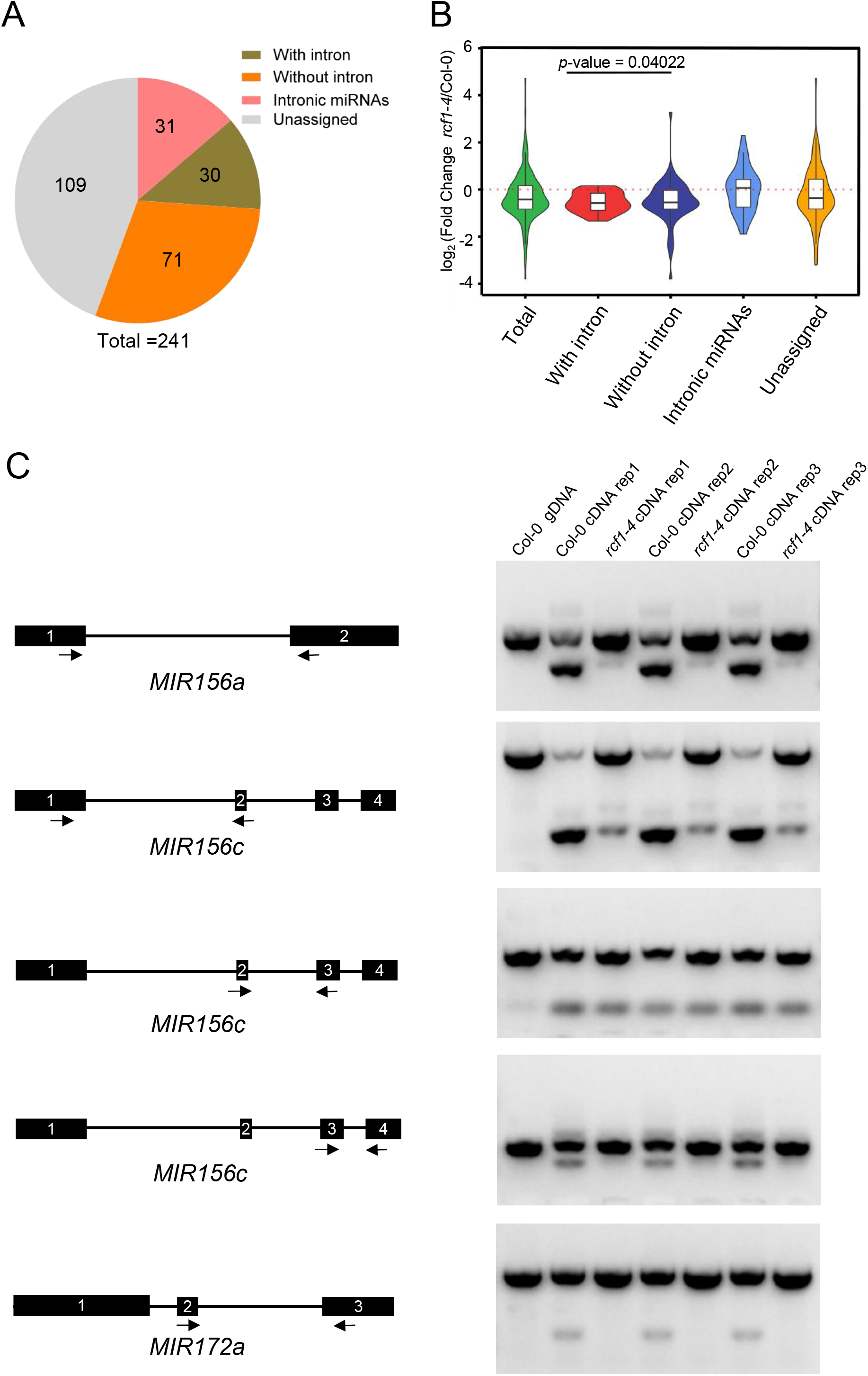
RCF1 affects the splicing of some pri-miRNAs. (A) 241 miRNAs detected in small RNA-seq were divided into 4 categories: with intron, without intron, intronic miRNAs and unclassified. (B) Violin plots showing the fold change of miRNAs in different categories in *rcf1-4* compared to Col-0. RT-PCR analysis of intron retention of pri-miRNAs. Black boxes and lines represent exons and introns respectively, arrows represent primers, the numbers in the rectangles indicate the exons of the *MIR* gene.

### RCF1 affects the splicing of pre-mRNAs

*Arabidopsis RCF1* and its homologous gene in rice *osRH42* were reported to be involved in pre-mRNA splicing under cold stress (Lu et al., 2020). To determine whether *RCF1* also affects pre-mRNA splicing under normal conditions, we performed poly A RNA-seq in Col-0 and *rcf1-4*. Clustering analysis show highly reproducible in the three biological replicates of RNA-seq in Col-0 and *rcf1-4*(Supplemental Figure S8). We analyzed the alternative splicing (AS) events in Col-0 and *rcf1-4*. Compared with wild type, thousands of differential splicing events were found in *rcf1-4*, including RI (retained introns), SE (skipped exons), A5SS (alternative 5’ splice sites), A3SS (alternative 3 ‘splice sites), and MXE (mutually exclusive exons) (Figure 7A; Supplemental Dataset S2). Genes with intron retention did not tend to be the differentially expressed genes in *rcf1-4*, indicating that RCF1-mediated alternative splicing did not affect the expression of these genes (Figure 7B; Supplemental Dataset S3). The alternative splicing events of some representative genes are shown in Figure 7C. For example, *AT5G13260* accumulated more exon-skipped transcripts; *AT3G01310* accumulated more transcripts with alternative 5’ splice sites; *AT2G37340* accumulated more transcripts with alternative 3’ splice sites; and *AT5G54750* accumulated more transcripts with mutually exclusive exons (Figure 7C). Interestingly, GO analysis of genes with AS events revealed that most of these differentially alternatively spliced genes were associated with RNA metabolism (Figure 7D; Supplemental Dataset S4). By IGV and RT-PCR analysis, we identified and validated the splicing changes in three RNA splicing-related genes (*RSZ33* (*AT2G37340*), RNA helicase family protein (*AT3G62310*) and *RS2Z32* (*AT3G53500*)) in *rcf1-4* (Figure 7E). Collectively, RCF1 is also involved in pre-mRNA splicing, which may affect miRNA biogenesis indirectly.

**Figure 7.**
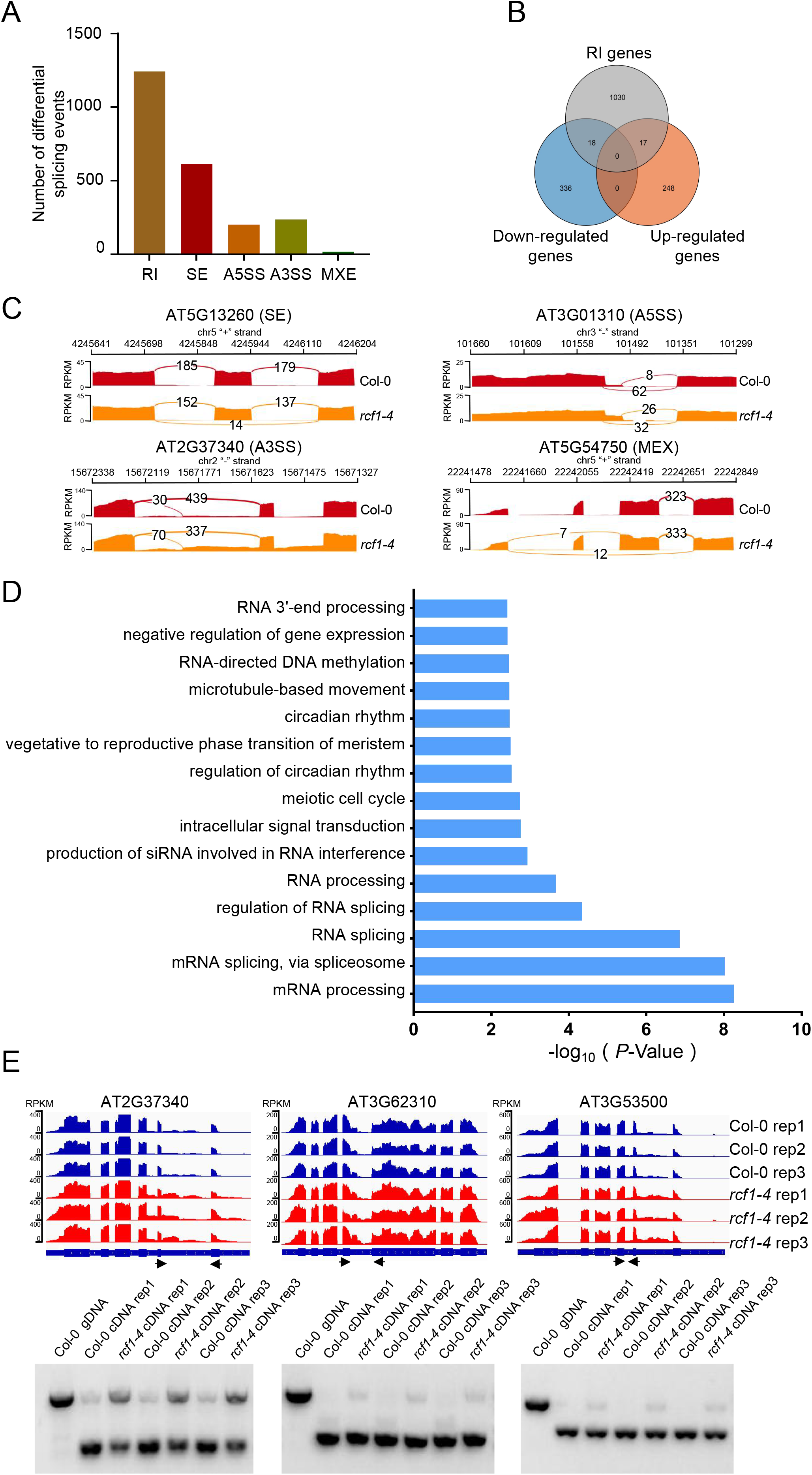
RCF1 is required in the splicing of pre-mRNAs. (A) Bar charts showing numbers of the significantly affected alternative splice sites between WT and *rcf1-4*. RI (retained introns); SE(skipped exons); A5SS(alternative 5′ splice sites); A3SS(alternative 3′ splice sites); MXE(mutually exclusive exons);. (B) Venn diagram showing the degree of overlap among RI genes, down-regulated genes and up-regulated genes in *rcf1-4*. (C) Examples of some AS events in *rcf1-4*. The y-axes are RPKM, the x-axis is the genomic position, the arcs indicate exon-exon junctions and the numbers are counts of exon-exon junctions. (D) Top 20 enriched Gene Ontology (GO) terms of DAS genes between WT and *rcf1-4*. (E) Examples of three genes related to pre-mRNA splicing with intron retention defects in WT and *rcf1-4*. Validation of affected IR events by RT-PCR analyses.

## Discussion

RCF1, a DEAD-box helicase, is conserved from yeast to mammals, (Xu et al., 2004). A previous study suggested that RCF1 is involved in pre-mRNA splicing under cold stress (Guan et al., 2013).However the biological functions of RCF1 under normal development conditions remain to be understood. In this work, we show that a point mutation in *RCF1* displays decreased accumulation of miRNAs globally, and the accumulation of several examined miRNA target transcripts are increased, indicating that the mutation in *RCF1* causes a defect in miRNA pathway. Consistent with the decrease in miRNA levels, the *rcf1-4* mutant displays pleiotropic developmental alterations.

MiRNA biogenesis initiates with the transcription of *MIR* genes. RCF1 does not enable miRNA biogenesis by promoting *MIR* transcription. The main pieces of evidence are: (1) pri-miRNA accumulation is increased rather than decreased in *rcf1-4*, and; (2) levels of *GUS* transcripts controlled by the *MIR167a* promoter were not affected in *rcf1-4*. The increased accumulation of pri-miRNAs but decreased accumulation of mature miRNAs in the *rcf1-4* mutant indicated that RCF1 promotes miRNA biogenesis at the pri-miRNA processing step. Proteins involved in *Arabidopsis* miRNA biogenesis can be divided into two groups based on their distinct effects on the accumulation of pri-miRNAs and mature miRNAs.

A group of proteins, for which mutants have reduced the accumulation of mature miRNAs and increased the accumulation of pri-miRNAs, promote the processing of pri-miRNAs. Including CBP80 and CBP20(Kim et al., 2008), SICKLE (Zhan et al., 2012), TOUGH (TGH) (Ren et al., 2012), STA1 (Ben et al., 2013), THO1 and THO2(Francisco-Mangilet et al., 2015), PSR1-INTERACTING PROTEIN 1 (PINP1)

(Qiao et al., 2015) and MOS2 (Wu et al., 2013). A second set of protein promote increased levels of mature miRNAs and pri-miRNAs. Representative proteins of this group include DAWDLE (DDL) (Yu et al., 2008), CDC5 (Zhang et al., 2013), Negative On TATA Less 2 (NOT2) (Wang et al., 2013), Elongator (Zhang et al.,2013), PRL1 (Zhang et al., 2014), MAC7 (Jia et al., 2017), SMALL 1 (SMA1) (Li et al., 2018), SEAP1 (Li et al., 2021), Reduction in Bleached Vein area (RBV) (Liang et al., 2022) and AAR2 (Fan et al., 2022). Our works indicate that RCF1 belongs to the first group of proteins that possibly promotes pri-miRNA processing.

RCF1 enables miRNA biogenesis by two mechanisms that may be related: D-body formation and the association of HYL1 with pri-miRNAs. Several miRNA biogenesis factors have been reported to be involved in D-body formation. Mutants of *MOS2* (Wu et al., 2013), *PRL1* (Zhang et al., 2014), *CDC5* (Zhang et al., 2013), *MAC7* (Jia et al., 2017), *PP4R3* (Wang et al., 2019), *THP1* (Zhang et al., 2020b), *RBV* (Liang et al., 2022), *AAR2* (Fan et al., 2022) and *SEAP1* (Li et al., 2021) exhibit a reduced number of D-bodies and show reduced levels of miRNAs. Similarly, D-body number and miRNA accumulation concomitantly decrease in the *rcf1-4* mutant. The accumulation of pri-miRNAs and HYL1 protein increased in the *rcf1-4* mutant compared with Col-0, suggesting that the impaired D-body formation is not caused by a reduction in these D-body constituents. So far little is known about the mechanism of D-body formation, but liquid-liquid phase separation may be a key factor in this process. Previous studies showed that AtRH6/AtRH8/AtRH12, three functionally redundant DEAD-box helicases, participate in miRNA biosynthesis by promoting D-body liquid-liquid phase separation in an RNA-and ATP-dependent manner in *Arabidopsis* (Li et al., 2021). Increasing studies have found that a common function of DEAD-box helicases is to regulate gene expression and various life processes by regulating the liquid-liquid phase separation of RNP condensates (Hondele et al., 2019; Weis and Hondele, 2022). It was reported that the IDRs (Intrinsically Disordered Regions) of AtRH6/8/10 and SE play a key role in regulating the liquid-liquid phase separation of D-bodies (Li et al., 2021; Xie et al., 2021). Similar to the structures of AtRH6/AtRH8/AtRH12, the N-terminal portion of the RCF1 protein is also essentially disordered. The ATPase activity of DEAD-box helicases is also important for its regulation of liquid-liquid phase separation (Hondele et al., 2019; Li et al., 2021; Weis and Hondele, 2022). In the *rcf1-4* mutant, the mutation site is located in the conserved ATP-binding region, and RCF1 with a mutation in the ATP-binding region cannot rescue the defects of *rcf1-1* (Guan et al., 2013), indicating that the ATPase activity of RCF1 is important for its function.

Some miRNA biogenesis factors enhance miRNA biogenesis by promoting the interaction between HYL1 with pri-miRNAs, such as MOS2 (Wu et al., 2013), DBR1 (Li et al., 2016) and STV1 (Li et al., 2017). RCF1 also promotes the binding of pri-miRNAs by HYL1, which was supported by the decreased association of HYL1 with pri-miRNAs in *rcf1-4,* despite the increased levels of pri-miRNAs and the HYL1 protein in *rcf1-4.* Given the smaller number of D-bodies in *rcf1-4*, it is possible that the reduced binding of pri-miRNAs by HYL1 in *rcf1-4* reflects reduced recruitment of pri-miRNAs into D-bodies.

As a potential core component of the spliceosome (Lu et al., 2020), RCF1 can promote the splicing of not only pre-mRNAs but also intron-containing pri-miRNAs, such as pri-miR156a, pri-miR156c, and pri-miR172a. The levels of the mature miRNAs processed from intron-containing pri-miRNAs are significantly lower than those from pri-miRNAs without introns in *rcf1-4*, suggesting that in addition to promoting D-body formation and pri-miRNA-HYL1 association, RCF1 enables the biogenesis of miRNAs processed from intron-containing pri-miRNAs by one additional mechanism, which is promoting the splicing of pri-miRNAs. RCF1-mediated pri-miRNA splicing probably works independently of RCF1-mediated pri-miRNA processing. The expression of intronic miRNAs is generally regulated by their host genes due to their special location (Yang et al., 2012). Through mRNA-seq, we found that the host genes containing intronic miRNAs are expressed at low levels in Col-0 and *rcf1-4* and there is no differential expression between these two genotypes (Supplemental Dataset S5). Although RCF1 affects the splicing of pre-mRNAs, the AS event did not impact the expression levels of the host genes. The intronic miRNAs are of very low abundance, which precluded determination of differential accumulation between *rcf1-4* and wild type.

Although both *OsRH42* and *AtRCF1* were reported to be induced by cold stress and participate in the splicing of pre-mRNAs under cold stress (Guan et al., 2013), a large number of AS events were also present in rice *OsRH42* mutants cultured under normal conditions (Lu et al., 2020). Unlike previous reports (Guan et al., 2013), we found that RCF1, as a homologous protein of OsRH42, also widely affects the splicing of pre-mRNAs at room temperature. Considerable differentially alternative splicing (DAS) genes involved in the regulation of RNA metabolism in the *rcf1-4* mutant, indicating that RCF1 may also be widely involved in RNA metabolism through direct and indirect manners.

Although dozens of proteins, such as SE, CBP20, CBP80 (Kim et al., 2008), CDC5 (Zhang et al., 2013), PRL1 (Zhang et al., 2014), MAC3A/MAC3B (Li et al., 2018), MAC7 (Jia et al., 2017), SMA1 (Li et al., 2018), THP1 (Zhang et al., 2020b), THO1, THO2 (Francisco-Mangilet et al., 2015), DBR1 (Li et al., 2016), HOS5 (Chen et al., 2015), PINP1 (Qiao et al., 2015), MOS2 (Wu et al., 2013), SCIKLE, STA1 (Ben et al., 2013), PP4 (Wang et al., 2019), RBV (Liang et al., 2022), AAR2 (Fan et al., 2022) and SEAP1 (Li et al., 2021) are involved in both pre-mRNA splicing and miRNA biogenesis events, these two processes can be independent. However, splicing defects in both pre-mRNAs and pri-miRNAs were only detected in mutants of CBP20 and CBP80 (Kim et al., 2008), HOS5 (Chen et al., 2015), STA1 (Ben et al., 2013), SEAP1 (Li et al., 2021). RCF1 is also involved in the splicing of both pre-mRNAs and pri-miRNAs. Perhaps RCF1 acts broadly in nuclear RNA metabolism, with miRNA biogenesis and RNA splicing being two independent processes that require RCF1.

## Supplemental Files

**Supplemental Figure S1.**
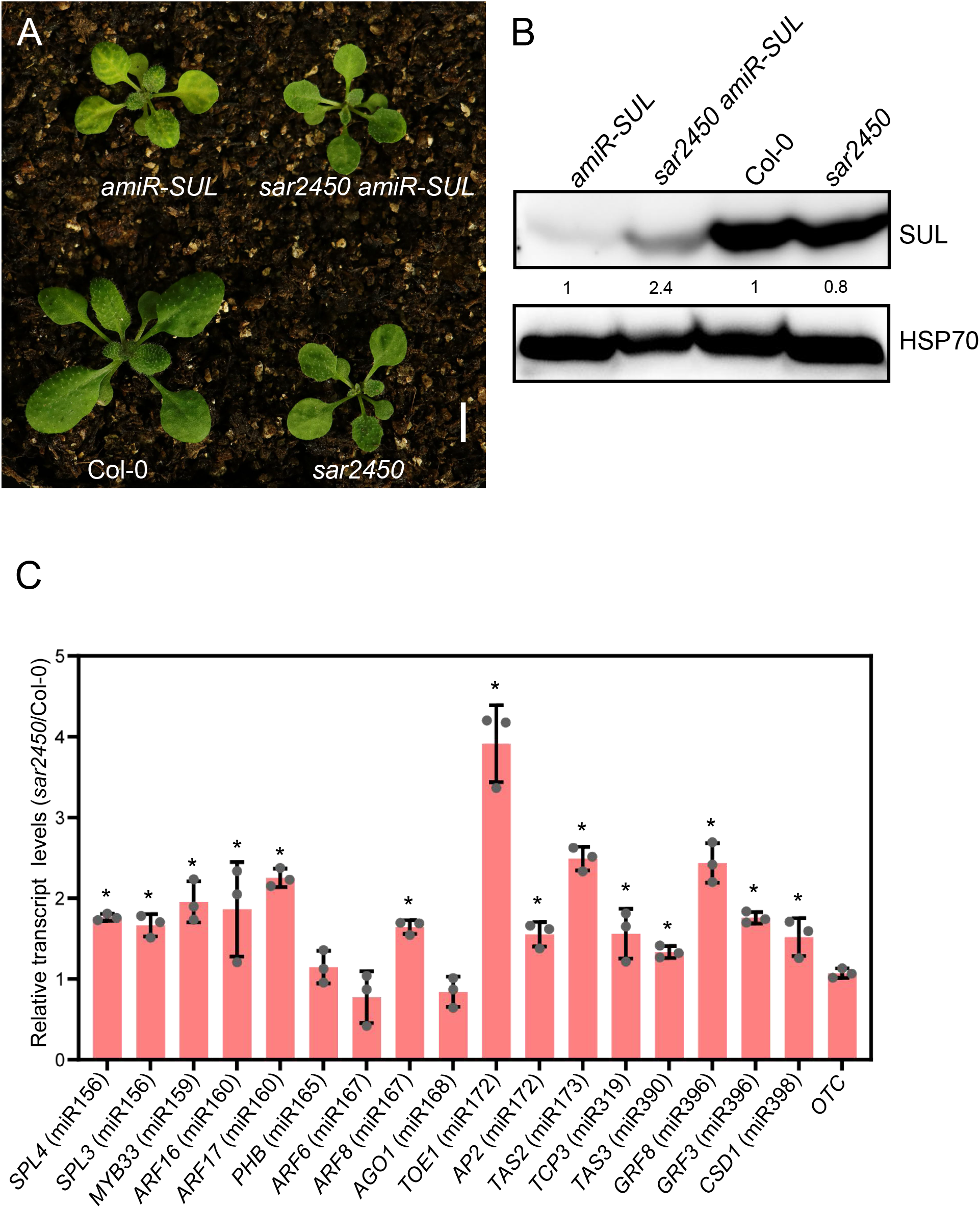
Comparison of the phenotypes and the expression of the target genes of amiR-SUL and some endogenous miRNAs between *sar2450* and the corresponding wild type. (A) Images of three-week-old plants of *amiR-SUL*, *sar2450 amiR-SUL*, Col-0 and *sar2450*. Bar = 1 cm. (B) Western blotting analysis of the SUL protein from *amiR-SUL*, *sar2450 amiR-SUL*, Col-0 and *sar2450* seedlings. The numbers represent relative abundance. (C) RT-qPCR analysis of miRNA target transcripts from *sar2450* and Col-0 seedlings. Transcript levels were normalized to those of *UBQ5* and compared with Col-0. *ORNITHINE CARBAMOYLTRANSFERASE* (*OTC*) is a housekeeping gene.(Error bars represent SD (standard deviation) of three replicates, *P < 0.05).

**Supplemental Figure S2.**
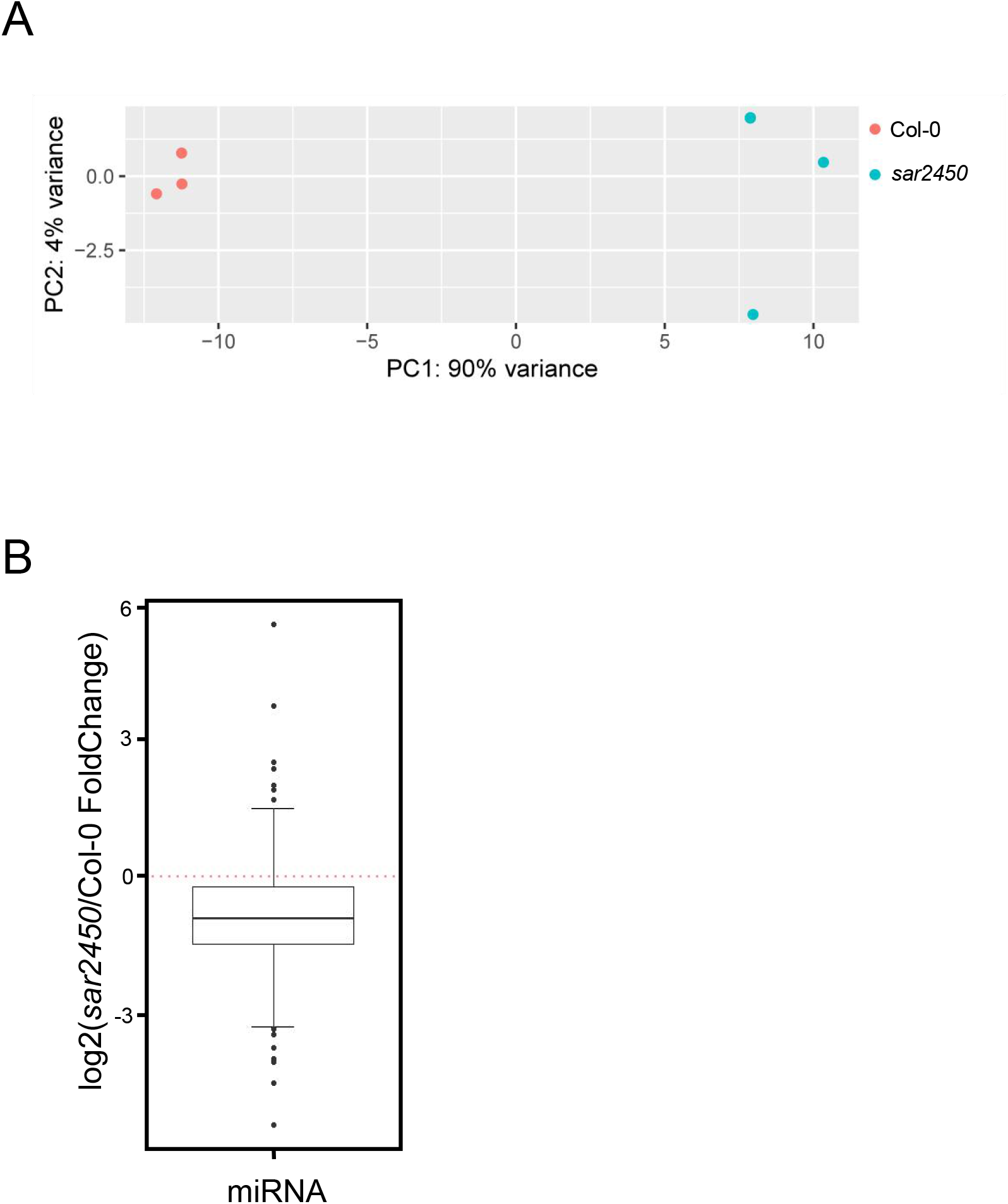
Supplemental Figure S2 The accumulation of miRNAs was globally reduced in *sar2450*. (A) Principal component analysis showed highly correlated in the three biological replicates of small RNA-seq in Col-0 and *rcf1-4*. (B) Box plots showing differentially accumulated miRNAs in WT and *sar2450* mutants.

**Supplemental Figure S3.**
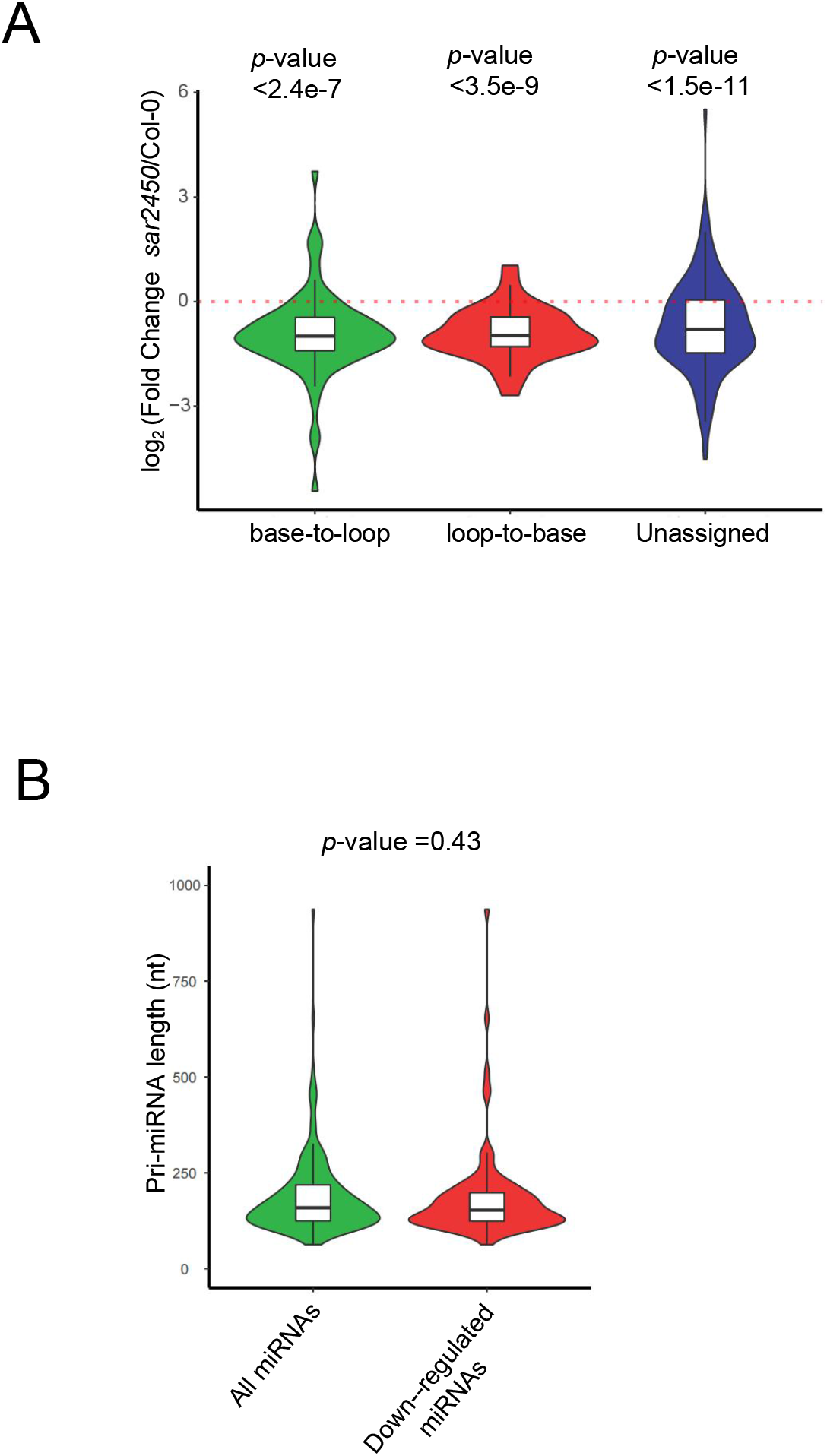
Violin and box plots showing the common features of down-regulated miRNAs in *sar2450*. (A) Differential accumulation of miRNAs with different processing directions in *sar2450* as compared with WT. (B) There was no significant difference in pri-miRNA length between all detected miRNAs and down-regulated miRNAs in *sar2450*.

**Supplemental Figure S4.**
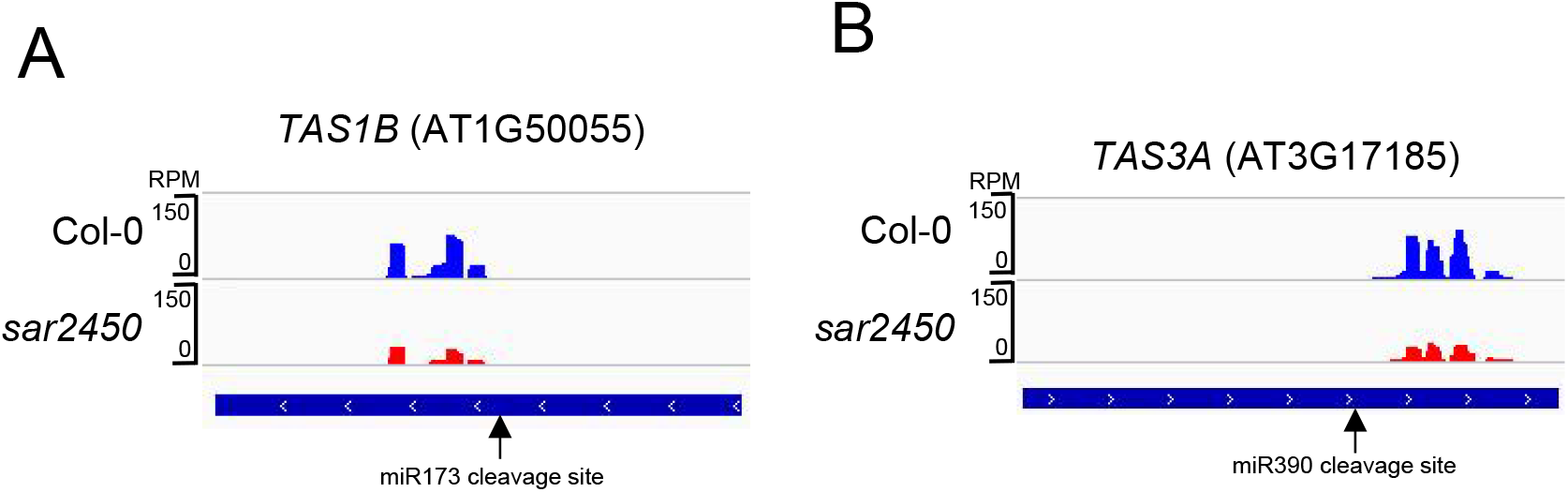
ta-siRNAs were down-regulated in *sar2450*. (A) and (B) IGV showing the accumulation of ta-siRNAs from *TAS1B* and *TAS3A* respectively was reduced in *sar2450*. Small RNA-seq reads are shown against the gene models below.

**Supplemental Figure S5.**
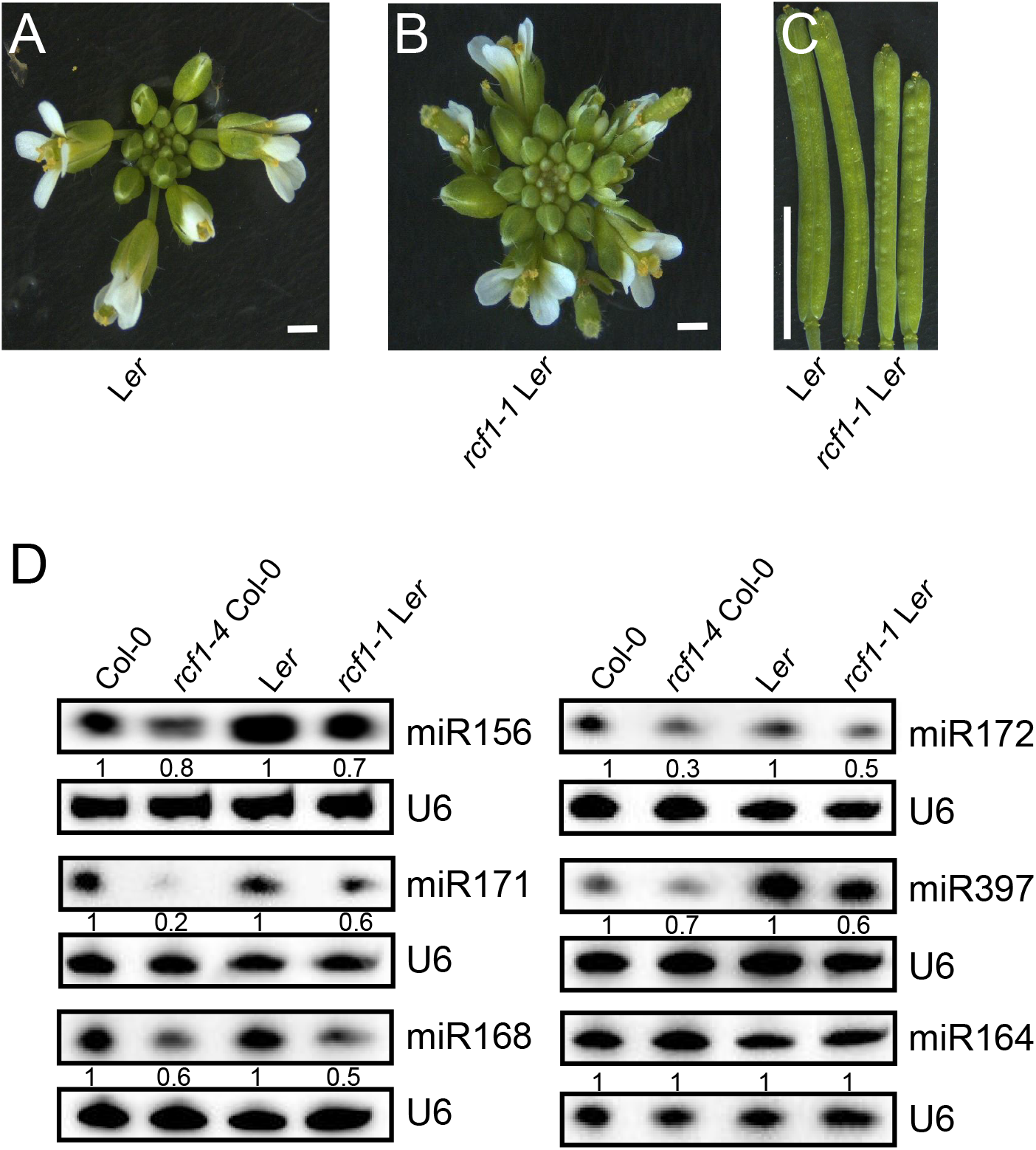
Mutant phenotypes and decreased miRNA accumulation of *rcf1-1*. (A-B) Images showing aberrant flowers and inflorescences in *rcf1-1*. Bar = 2 mm. (C) Images of mature siliques showing reduced silique lengths in *rcf1-1*. Bar = 1 cm. (D) Northern blotting analysis of miRNAs in Col-0, *rcf1-4* (in Col-0), L*er* and *rcf1-1* (in L*er*). U6 RNA served as a loading control. The numbers below the gel images represent the relative amount.

**Supplemental Figure S6.**
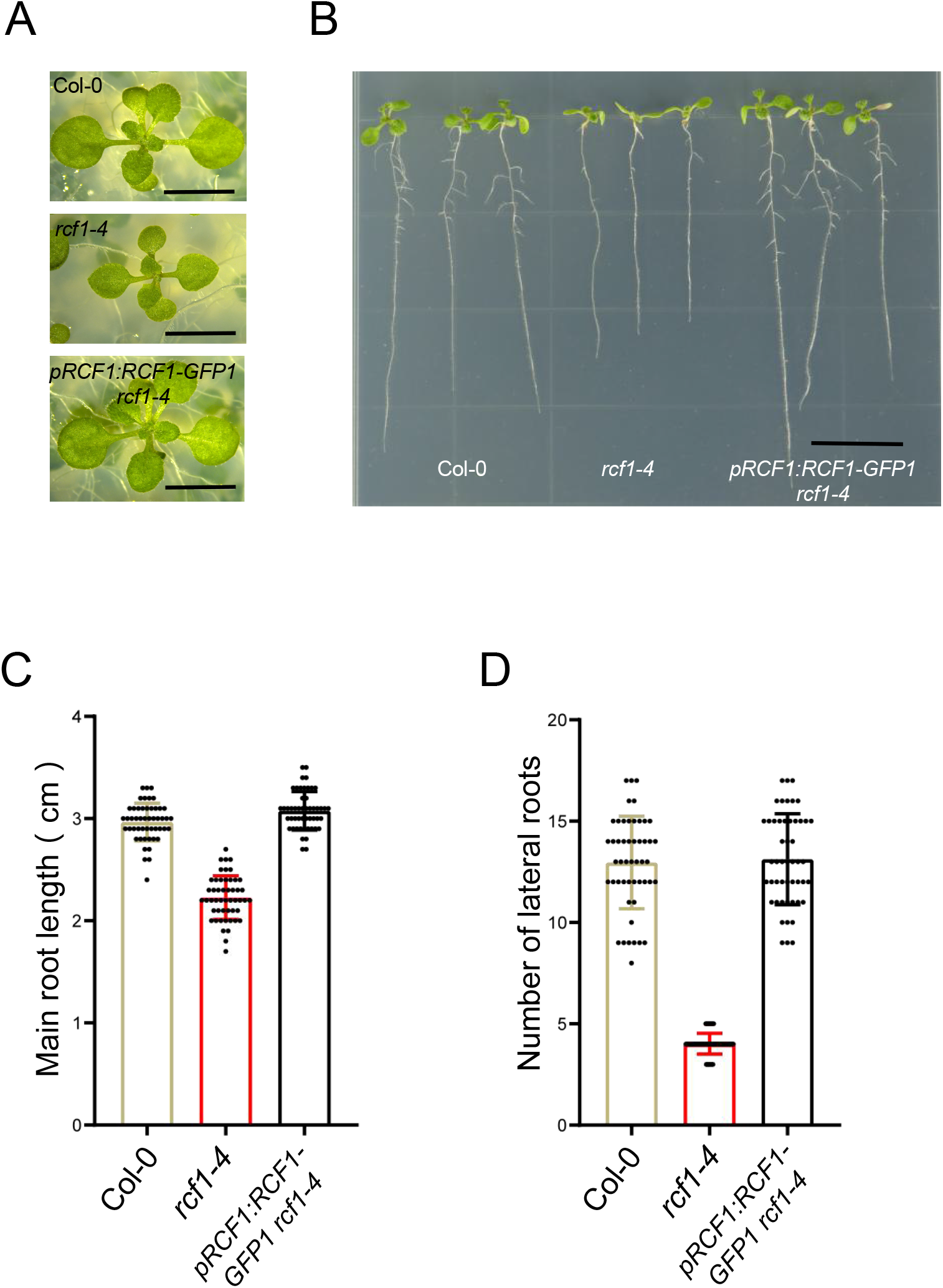
The *pRCF1:RCF1* transgene restores the *rcf1-4* mutant phenotypes. (A) 14-day-old seedlings of Col-0, *rcf1-4*, and *pRCF1:RCF1-GFP1 rcf1-4*. Bar = 1 cm. (B) Images of 10-d-old seedlings of the genotypes as indicated. Bar = 1 cm. (B) Measurements of the main root length of 10-day-old seedlings of the genotypes as indicated. Error bars represent SD (n = 50). (C) Number of lateral roots of 10-d-old seedlings of the genotypes as indicated. Error bars represent SD (n = 50).

**Supplemental Figure S7.**
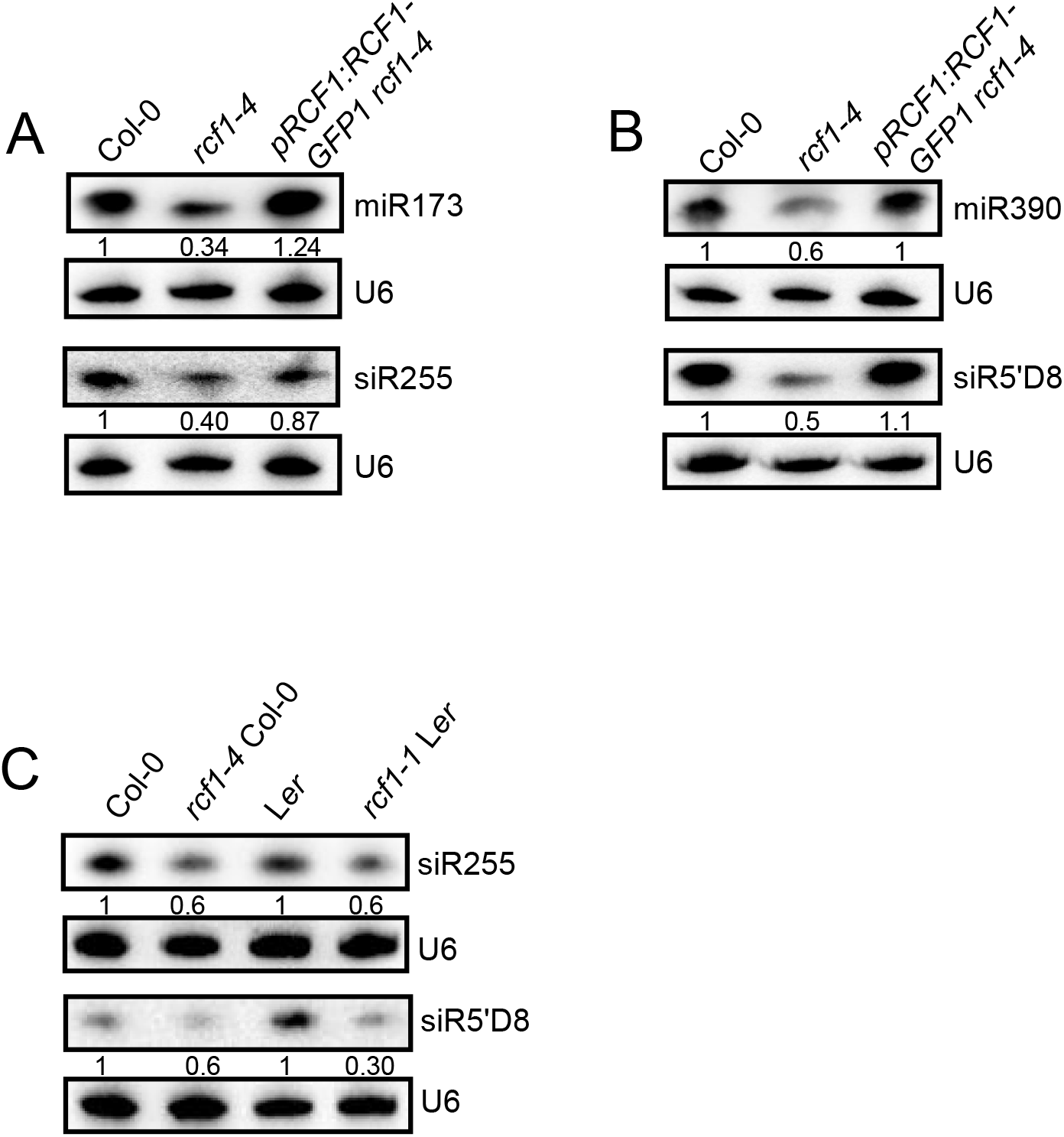
RCF1 promotes the accumulation of ta-siRNAs. (A-B) Northern blotting analysis showing of ta-siRNAs and the corresponding trigger miRNAs in Col-0, *rcf1-4*, and *pRCF1:RCF1-GFP1 rcf1-4*. U6 was used as a loading control. The numbers below the gel images represent the relative amount. (C-D) Northern blotting analysis of ta-siRNAs in Col-0, *rcf1-4* (in Col-0), L*er* and *rcf1-1* (in L*er*). U6 was used as a loading control. The numbers below the gel images represent the relative amount.

**Supplemental Figure S8.**
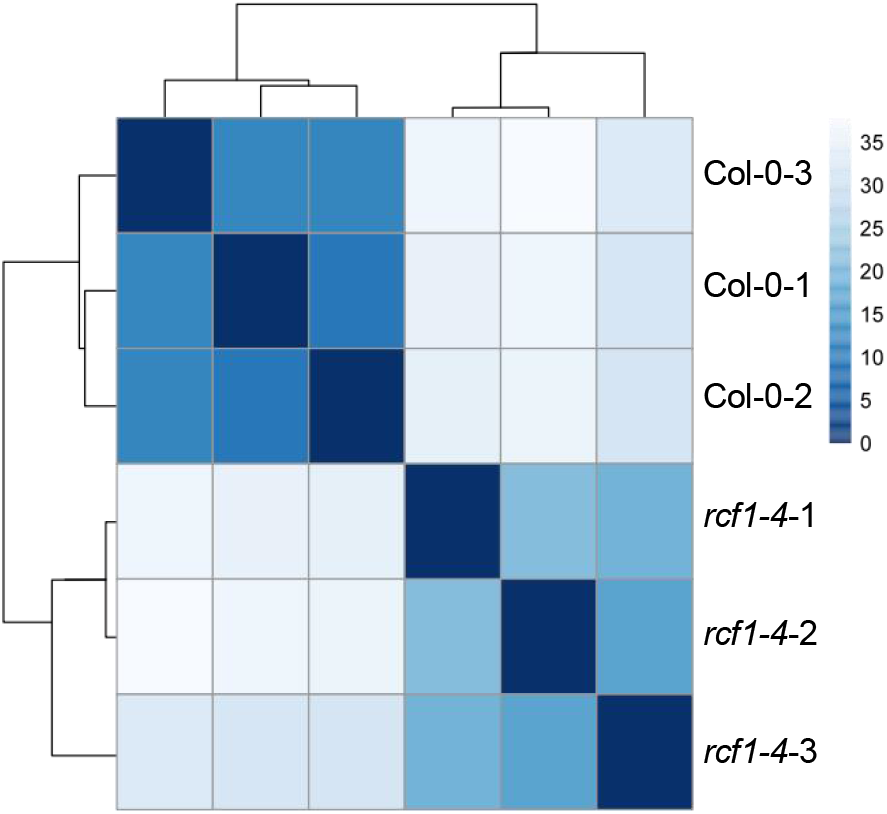
Clustering analysis showed highly correlated in the three biological replicates of RNA-seq in Col-0 and *rcf1-4*.

Supplemental Dataset S1. miRNAs from small RNA sequencing of Col-0 and *rcf1-4*. Supplemental Dataset S2. The lists of DAS genes between WT and *rcf1-4*.

Supplemental Dataset S3. Differentially expressed genes identified between Col-0 and *rcf1-4*.

Supplemental Dataset S4. GO term enrichment results of DAS genes between WT and *rcf1-4*.

Supplemental Dataset S5. The expressing of host genes containing intronic miRNAs in Col-0 and *rcf1-4*.

Supplemental Dataset S6. Primers and probe sequences.

## Methods

### Plant Materials and Growth Conditions

*rcf1-4 amiR-SUL* is a novel point mutant isolated from *pSUC2:amiR-SUL* mutagenesis screening. The *rcf1-1* (in L*er* background) is a kind gift from Jianhua Zhu. Transgenic lines *pSUC2:amiR-SUL*, *pHYL1:HYL1-YFP*, and *pMIR167a:GUS* were previously described(Liang *et al*., 2022). For plate-cultured seedlings, all seeds were sewn on 1/2 Murashige and Skoog Basal Medium containing 1% Sucrose and 0.8% Agar. Plants were grown at 22 ^°^C under 16 h light: 8 hdark cycles.

### Genetic Mapping

Genetic mapping was conducted as described(Liang et al., 2022). In the F2 population, pooled genomic DNA of ∼200 plants with the *sar2450 amiR-SUL* phenotype, and subjected to library construction. The library was pair-end sequenced on the Illumina platform Hiseq2000 at 50× coverage at BGI-Shenzhen, China. The mutation in AT1G20920 (*RCF1*) was confirmed by sequencesing the PCR products amplified from *RCF1*.

### DNA constructs and complementation

The genomic fragment of *RCF1* (At1g20920) including ∼2.5 kb promoter and the coding region without the stop codon was PCR amplified from Col-0 genomic DNA with primers RCF1-F and RCF1-R (Supplemental Dataset S6) and cloned into the pTSK108 vector. The vector was recombined with pMDC107(Curtis and Grossniklaus, 2003) via LR recombination to generate the pRCF1:RCF1-GFP construct. The *pRCF1:RCF1-GFP* construct was introduced into *rcf1-4 amiR-SUL*. The transgenic plants were identified by hygromycin resistance screening.

### Fluorescence microscopy

For the visualization of HYL1-YFP fluorescence in *pHYL1:HYL1-YFP* and *pHYL1:HYL1-YFP rcf1-4*, roots of seedlings grown at 1/2 Murashige and Skoog Basal Medium containing were observed using a LeicaSP8 confocal microscope (excitation, 488nm; emission, 520 nm).

### Histochemical GUS staining

For visualize GUS expression, whole seedlings from *pMIR167a:GUS* and *pMIR167a:GUS rcf1-4* were immersed in the GUS staining solution for 30 minutes at 37 °C in the dark. Stained seedlings were cleared with 70% ethanol to remove chlorophyll before imaging under a stereomicroscope (Leica).

### RT-qPCR

Total RNAs from inflorescences or seedlings were treated with DNase I followed by reverse transcription using PrimeScript™ II 1st Strand cDNA Synthesis Kit (TAKARA, 6210A) with oligo-d(T) primers. RT-qPCR was performed using the SYBR premix ExTaq II kit (TAKARA, RR820A) on the BioRad CFX96 system. The primers for target genes and pri-miRNAs are listed in Supplemental Dataset S6.

### Small RNA sequencing and data analysis

Total RNA was extracted from 14-day-old seedlings. Twenty micrograms of total RNA were subjected to 15% urea-denatured polyacrylamide gel electrophoresis, and small RNAs of 15-40 nt were excised from the gel and used to construct a library using “NEB Next Multiplex Small RNA Library Prep Set” for Illumina (New England Biolabs, E7300). High-throughput sequencing using the Illumina NextSeq500 platform was performed at BerryGenomics, China. Sequencing data were analyzed by pRNASeqTools (https://github.com/grubbybio/pRNASeqTools). Normalization was performed by calculating the RPM value (reads per million trimmed reads).

### Northern blot analysis

Northern blotting was performed as described(Liang et al., 2022). Total RNA was extracted from 14-day-old seedlings, and 5 μg of total RNA was separated by 15% (wt/vol) acrylamide/7 M urea gel and subsequently transferred to a Hybond-NX nylon membrane (GE healthcare) after gel electrophoresis. The 5’ and 3’ ends of antisense complementary oligonucleotides probes were modified with biotin. Membranes and probes were hybridized overnight at 55°C. The biotin signals on the membrane were detected using the chemiluminescent nucleic acid detection module (Thermo Fisher, 89880).

### Western blot analysis

Western blots were performed as described(Liang et al., 2022). 0.1 g of 14-day-old seedlings were ground into powder in liquid nitrogen and proteins were then extracted with protein extract solution (50 mM Tris-HCl pH 8, 150 mM NaCl, 0.1% SDS, 1 mM EDTA, 0.1% sodium deoxycholate, 1% Triton X-100, 5 mM DTT, and 1% proteinase inhibitor cocktail). Total proteins were separated by 12% (v/v) SDS polyacrylamide gel electrophoresis and transferred to Hybond C-Extra membranes (Amersham Biosciences). Antibodies used in this study included anti-HSP70 (Agrisera, AS08371; dilution, 1:10000), anti-SUL (dilution, 1:2000)(Jia et al., 2017), anti-HYL1 (Agrisera, AS06136; dilution, 1:2000), anti-SE (Agrisera, AS09532A; dilution, 1:1000), and anti-histone H3 (Agrisera, AS10710; dilution 1:3000). To generate anti-DCL1 antibodies, the purified 1-250 amino acids of DCL1 was used as antigens to raise polyclonal antibodies in rabbits at ABclonal Technology Co.Ltd.

### RNA-Seq data analysis

Total RNA was extracted from inflorescences, and mRNA was isolated using VAHTS mRNA Capture Beads (Vazyme, N401-01). The mRNA library was constructed and sequenced at Novogene, China. DEGs were identified between Col-0 and *rcf1-4* by pRNASeqTools (https://github.com/grubbybio/pRNASeqTools) with fold-change >2 < 0.01 and *P*-value as filters. Alternative splicing (AS) events in Col-0 and *rcf1-4* were analyzed with rMATS (Shen *et al*., 2014) and using FDR < 0.1, Diff_PI_Density > 0.05 as filters.

### RIP

Four grams of 14-day-old Col-0 or *rcf1-4* seedlings were harvested and cross-linked with 1% formaldehyde (1% formaldehyde, 1 mM PMSF, 10 mM Tris–HCl pH 8, 1 mM EDTA, and 0.4 M sucrose). The cross-linked tissue was ground into powder in liquid nitrogen and resuspended in Honda Buffer (1.25% Ficoll, 0.44 M sucrose, 2.5% Dextran T40, 10 mM MgCl2, 20 mM Hepes KOH pH 7.4, 0.5% Triton X-100, 1 mM PMSF, 5 mM DTT, and 1% proteinase inhibitor cocktail) with 8 U/mL RNase inhibitor. The resuspended slurry was filtered through two layers of Miracloth and centrifuged at 3000g for 7.5 minutes at 4°C. The pelleted nuclei were resuspended in nuclei lysis buffer (50 mM Tris-HCl pH 8, 10 mM EDTA, 0.1% SDS, 1% Triton X-100, 0.1% sodium deoxycholate, 150 mM NaCl, 5 mM DTT, and 1% proteinase inhibitor cocktail) with 160 U/mL RNase inhibitor and the nuclei were disrupted by sonication (Covaris S200). The sonicated samples were centrifuged at 13,800g for 5 minutes at 4°C to pellet debris. The supernatant was precleared with protein A agarose beads and then incubated with anti-HYL1 antibodies conjugated to protein A agarose beads with gentle rotation for 4 hours at 4°C. The beads were washed five times with nuclei lysis buffer with 40 U/mL RNase inhibitor and once with LiCl wash buffer (0.25 M LiCl, 1% sodium deoxycholate, 10 mM Tris-HCl pH 8.0, 1% CA630, and 1 mM EDTA) with 40 U/mL RNase inhibitor, and twice with TE buffer (10 mM Tris-HCl pH 8.0, 1 mM EDTA) with 40 U/mL RNase inhibitor. The immuno-complexes were eluted with elution buffer (1% SDS, 100 mM NaHCO3) at 65°C. Then, 200 mM NaCl and proteinase K were added to the immuno-complexes to reverse the crosslinking at 65°C for 12 hours. RNAs were extracted by Trizol and used as templates for RT-qPCR analyses, using primers listed in (Supplemental Dataset S6).

## Author Contributions

Chi Xu, Beixin Mo and Xuemei Chen conceived and designed the experiments. Chi Xu, Zhanhui Zhang, Juan He and Yongsheng Bai performed the experiments. Chi Xu, Zhanhui Zhang and Beixin Mo wrote the article. Xuemei Chen, Jihua Tang, Guiliang Tang and Lin Liu revised the article. All authors read and approved the final manuscript.

## Acknowledgments

We thank Dr. Guan for the gift of *rcf1-1*seeds. This research was funded by National Natural Science Foundation of China (No. 32270595, No. 32171985).

## Data availability

All raw and processed data were deposited at NCBI (http://www.ncbi.nlm.nih.gov/sra) under the accession number PRJNA948910. Source data are provided with this paper.

